# Visualizing the Geography of Genetic Variants

**DOI:** 10.1101/068536

**Authors:** Joseph H. Marcus, John Novembre

**Author notes:** Address correspondence to JHM or JN.

## Abstract

One of the key characteristics of any genetic variant is its geographic distribution. The geographic distribution can shed light on where an allele first arose, what populations it has spread to, and in turn on how migration, genetic drift, and natural selection have acted. The distribution of a genetic variant can also be of great utility for medical/clinical geneticists. Collectively the geographic distribution of many genetic variants can reveal population structure. As a result, visual inspection of geographic maps for genetic variants is common practice in genetic studies. Here we develop an interactive visualization tool for rapidly displaying the geographic distribution of genetic variants. Through a REST API and dynamic front-end the *Geography of Genetic Variants (GGV)* browser provides maps of allele frequencies in populations distributed across the globe.

## Introduction

Genetics researchers often face the problem that they have identified one or many genetic variants of interest using an approach such as a genome-wide association study and then would like to know the geographic distribution of the variant. For example, the researcher may hope to address: 1) implications for genomic medicine (e.g. Is a risk allele geographically localized to a certain patient population? What population should be studied to observe variant carriers? ROSENBERG *et al.*, 2010); or 2) the evolutionary history of the variant in question (e.g. does the variant correlate with a known environmental factor in a manner suggestive of some geographically localized selection pressure? NOVEMBRE and DI RIENZO, 2009; COOP *et al.*, 2010). A simple geographic map of the distribution of a genetic variant can be incredibly insightful for these questions.

Contemporary population genetics researchers are also faced with the challenge of large, high-dimensional datasets. For example, it is not uncommon for a researcher in human genetics to have a dataset comprised of thousands of individuals measured at hundreds of thousands or even millions of single nucleotide variants (SNVs). One common approach to visualizing such high-dimensional data is to compress the SNV dimensions down to a small number of latent factors, using a method such as principal components analysis (PRICE *et al.*,2006; PATTERSON *et al.*, 2006), or a model-based clustering method such as STRUCTURE (PRITCHARD *et al.*, 2000) or ADMIXTURE (ALEXANDER *et al.*, 2009). While these approaches are extremely valuable, researchers can use them too often without inspecting the underlying variant data in more detail. A natural approach to gaining more insight to the overall structure of a population genetic dataset is to visually inspect what geographic patterns arise in allele frequency maps.

Unfortunately, generating geographic allele frequency maps is time-consuming for the average researcher as it requires a combination of data-wrangling methods (KANDEL *et al.*, 2011) and map-making techniques that are unfamiliar to most. Our aim here is to produce a tailored system for rapidly constructing informative geographic maps of allele frequency variation.

Our work is inspired by past tools such as the ALFRED database (RAJEEVAN *et al.*, 2012) and the maps available on the HGDP Selection browser (PICKRELL *et al.*, 2009). One of us (JN) developed the scripts for the HGDP Selection Browser maps using The Generic Mapping Tools (GMT) (WESSEL *et al.*, 2013), a powerful system of geographic plotting scripts for making static plots. The plots from the HGDP Selection Browser have proved useful, have appeared in research articles (e.g. PICKRELL *et al.*, 2009; COOP *et al.*,2009), books (e.g. DUDLEY and KARCZEWSKI, 2013), and have been made available on the UCSC Genome Browser (available under the HGDP Allele Freq track of the browser KENT *et al.*, 2002).

Reference datasets for population genetic variation have greatly expanded since the release of the HGDP Illumina 650Y dataset (LI *et al.*, 2008) that formed the basis of the HGDP Selection Browser maps. The most notable advance is the publication of the 1000 Genomes Phase 3 data (THE 1000 GENOMES PROJECT CONSORTIUM, 2015) though additional datasets are continually coming online (e.g. LAZARIDIS *et al.*, 2014). In addition, novel approaches for data visualization have become more widely available. In particular, web-based visualization tools, such as Data Driven Documents (D3.js), offer useful methods for interactivity, the advantages of software development in modern web-browsers, a large open-source development community, and ease of sharing (BOSTOCK *et al.*, 2011).

Taking advantage of these recent advances, we aim to address the significant visualization challenges that are inherit in the production of geographic allele frequency maps, including dynamic interaction, display of rare genetic variation, and representation of uncertainty in estimated allele frequencies due to variable sample sizes.

## Fundamental Approach

The Geography of Genetic Variants browser (GGV) uses the scalable vector graphics and mapping utilities of D3.js (BOSTOCK *et al.*, 2011) to generate interactive frequency maps, allowing for quick and dynamic displays of the geographic distribution of a genetic variant. The front-end provides legends for the map and various configuration boxes to allow users to query different datasets or choose visualization options.

In order to allow for easy access to commonly used public genomic datasets, such as the 1000 Genomes project (THE 1000 GENOMES PROJECT CONSORTIUM, 2015) or Human Genome Diversity project (LI *et al.*,2008), we have developed a REST API (GRINBERG, 2014) for accessing data. The API allows querying of allele frequencies by chromosome and position, by reference SNP identifier (SHERRY *et al.*, 2001), or randomly sampled SNPs. While many applications require inspection of the distribution of a specific variant, from our experience, it can be very helpful to view the geographic distribution of several randomly chosen variants to quickly gain a sense of structure in a dataset. We find this to be especially useful in teaching contexts, as it provides a highly visual way for learners to understand human genetic variation.

After a query, the GGV displays the allele frequencies for a set of populations as a collection of pie charts where each represents the minor and major allele frequency in a single population. Pie charts are displayed as points at a latitude and longitude assoicated with a population and the map boundaries are chosen based off of the geographic configuration of populations in a given dataset [Figure 1].

**Figure 1:**
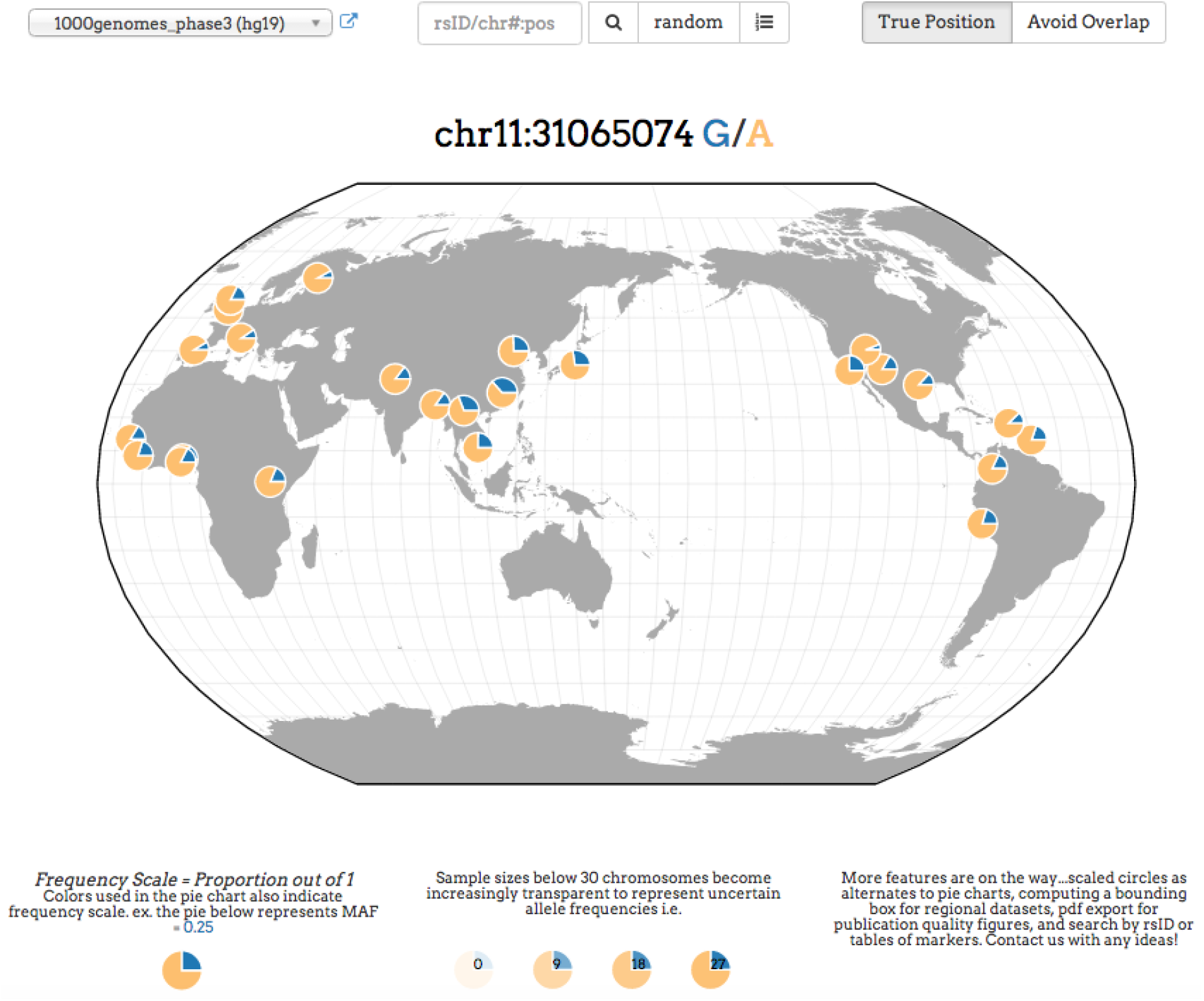
Example screenshot from the Geography of Genetic Variants browser using THE 1000 GENOMES PROJECT CONSORTIUM (2015) data. Each pie chart represents a population with the blue slice of the pie displaying the frequency of the global minor allele and the yellow slice of the pie displaying the frequency of the global major allele in each population.

## Representing uncertainty in frequency data

One under-appreciated problem with allele frequency maps is that not all data points have equal levels of certainty. For some locations, sample sizes are small, and the reported allele frequency may be quite far from the true population frequency due to sampling error. To address this issue, we use varying transparency in a population's pie chart: estimated frequencies with higher levels of sampling error (e.g. those from samples with *n* < 30) are made more transparent, and hence less visible, on the map [Figure 2].

**Figure 2:**
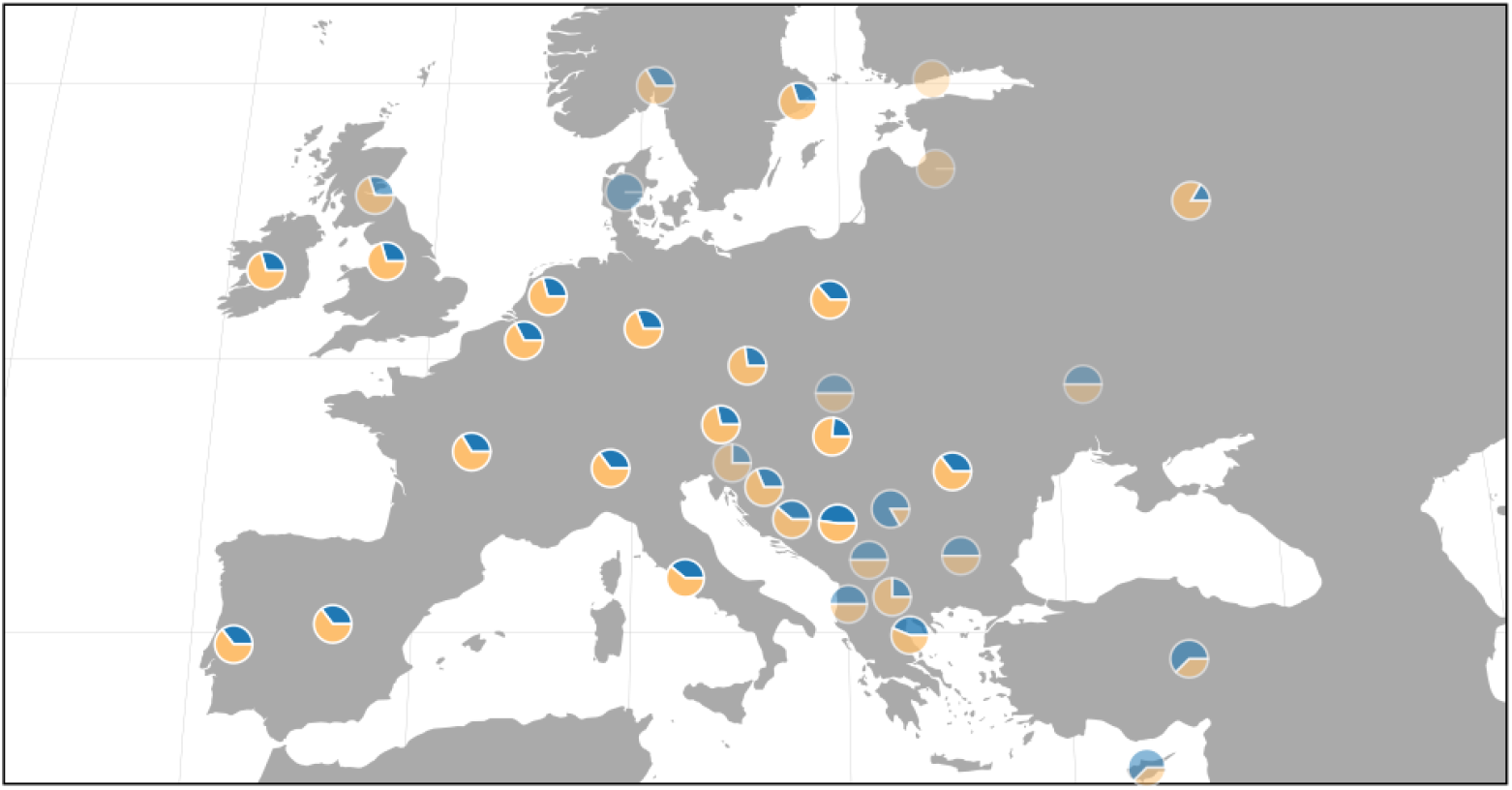
Example map from the Geography of Genetic Variants browser displaying the use of varying transparency of population pie charts to represent uncertainty in allele frequencies. The transparency is scaled in proportion to the number of observed chromosomes in each population for a particular variant. The frequency data and population identifiers are from NOVEMBRE *et al.* (2008).

## Representing rare variants in frequency data

An additional challenge is that allele frequencies between variants often differ greatly, sometimes by orders of magnitude in a single dataset. This has not been a pervasive problem until recently, as most population genetic samples were genotyped on SNP arrays, which have been biased towards variants that are common in human populations (5-50 % in minor allele frequency). With the combination of next generation sequencing technologies, new array designs focusing on rarer variants, and studies with thousands of individuals or more, it is now routine for the majority of variants to be rare (e.g. THE 1000 GENOMES PROJECT CONSORTIUM, 2015; NELSON *et al.*, 2012; TENNESSEN *et al.*, 2012). In visualization schemes using proportional area to represent frequency (such as standard pie charts), rare variants would be represented as narrow slivers, nearly invisible to the naked eye.

To address this challenge we re-scale frequencies for rare variants, so that small frequencies become visible. Specifically, we use a frequency scale that is indicated in a legend below the map and represented by varying color in the pie charts [Figure 3]. Much like scientific notation, this allows a wide range of frequencies to be displayed (Table 1).

**Figure 3:**
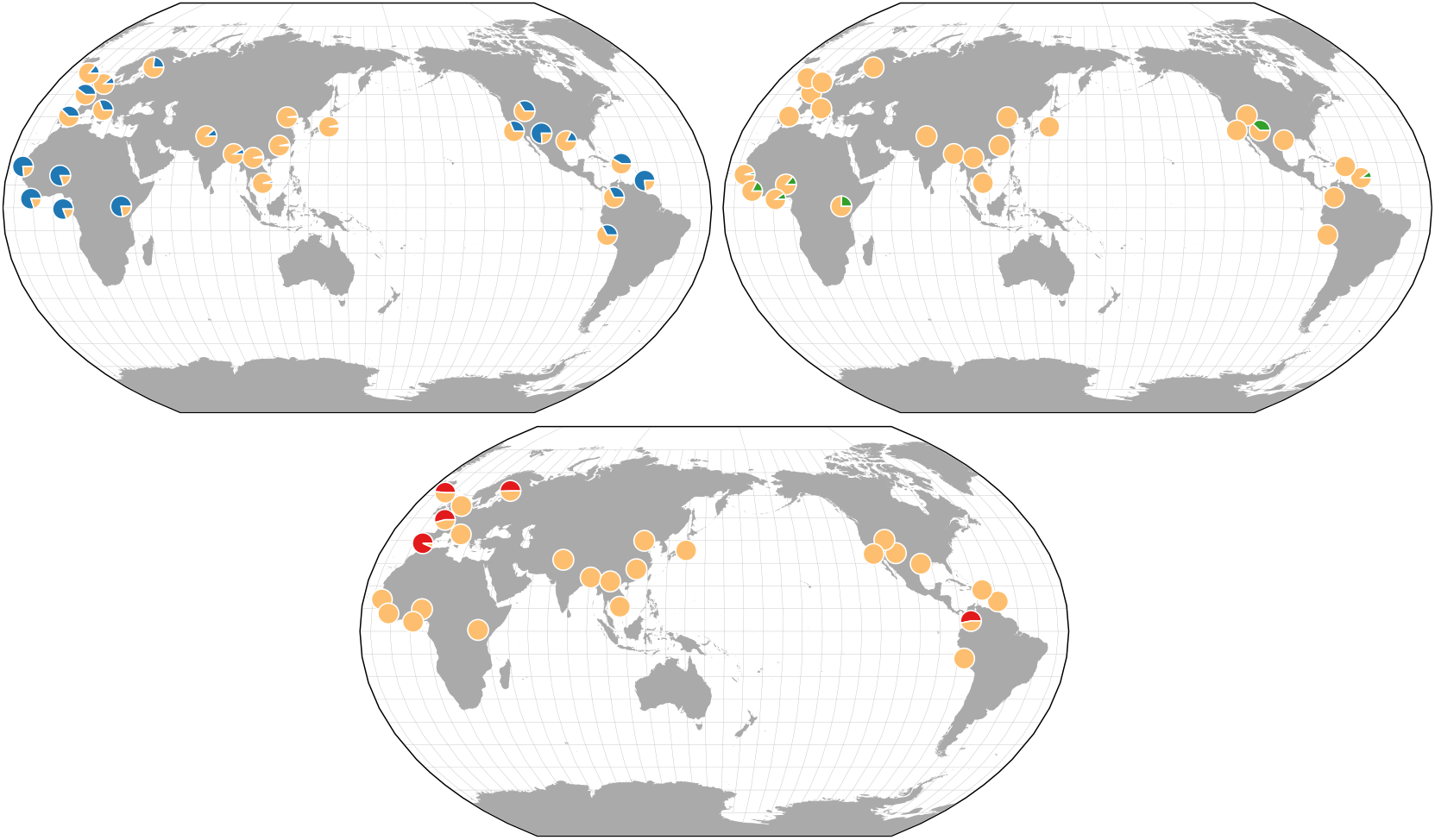
Example maps from the Geography of Genetic Variants browser displaying the use of frequency scales for more expressive representations of rare variation on geographic maps. The blue pie charts convey a given minor allele frequency out of 100 percent, the green out of 1 percent, and the red out of 0.01 percent. The data are from THE 1000 GENOMES PROJECT CONSORTIUM (2015).

**Figure 4:**
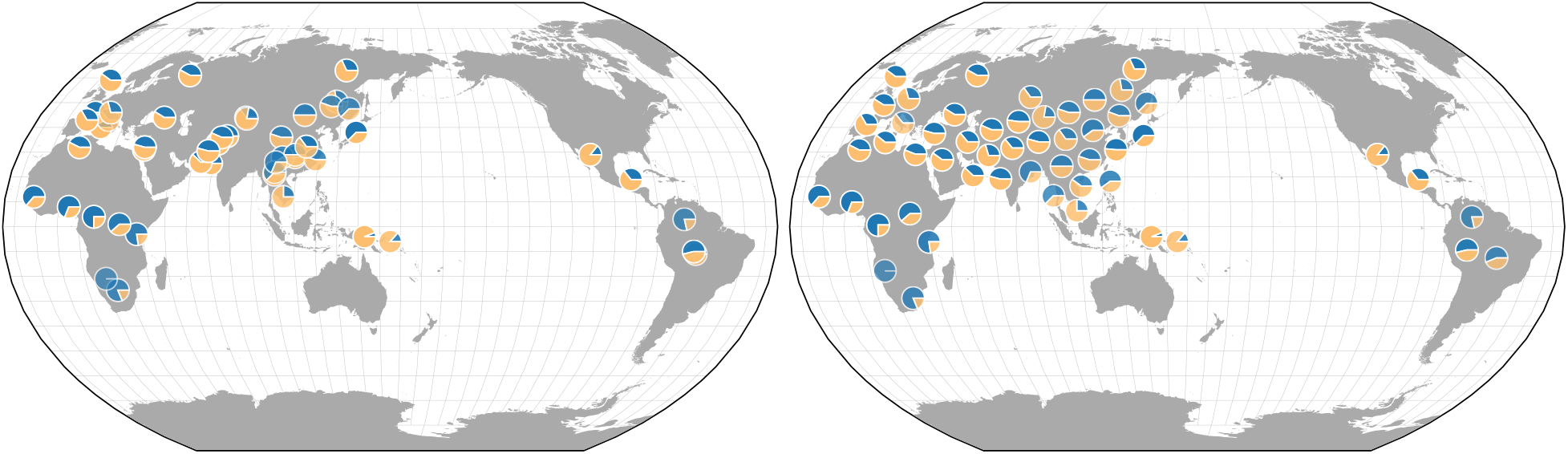
Example maps from the Geography of Genetic Variants browser displaying the use of a force directed layout to limit visual clutter when many populations overlap in geographic position. The left map shows the original population locations while the right shows the application of the force directed layout.

**Table 1:** Rare variants present a challenge for display. To address this challenge, the GGV browser changes the displayed image and the frequency scale of the map depending on the input sample frequency. As an example, a variant with a frequency 0.0025 is shown as a pie-chart that is 25% full and a frequency scale of 0.01 is marked in the legend of the map.

| Sample Frequency | Display Frequency Scale | Displayed Image |
| --- | --- | --- |
| 0.25 | 1 |  |
| 0.025 | 0.1 |  |
| 0.0025 | 0.01 |  |
| 0.00025 | 0.001 |  |

## Additional features of the interface

In many datasets where populations are sampled densely in geographic space, one problem is that allele frequency plots begin to overlap each other and obscure information. To address this issue, we use force-directed layouts of the populations such that no two points are overlapping each other, and yet the points will be pulled towards their true origins [Figure 5]. Also, by hovering the mouse cursor over any population, a user can see the population labels and precise frequency information.

## Access to the underlying frequency data

To provide an interface to the population minor allele frequency data, we use a REST API implemented in the python library Flask-RESTful (GRINBERG, 2014). The front-end D3.js visualization uses the API to obtain the data, though users can also interface with it directly. For the front-end, HTTP GET requests return json formatted allele frequency data and the meta-data associated with each population and genetic variant (e.g. latitude, longitude, population label, sample size, and frequency scale). Genetic variants can be queried by chromosome position, rsid, or randomly. Example HTTP requests and json response can be seen in the Appendix.

## Discussion

By allowing rapid generation of allele frequency maps, we hope to facilitate the interpretation of variant function and history by practicing geneticists. We also hope the ability to query random variants from major human population genetic samples will allow students to appreciate the structure of human genetic diversity in a more approachable and intuitive form than alternative visualizations.

A major challenge of using a geographic representation of genetic variation in humans is that the samples must be associated with a geographic location. While doing so is generally immensely helpful, it has inherent complexity and limitations. For example, practitioners must make choices regarding representing where an individual was sampled for the study (e.g. the city of a major research center) or choosing a location that is more representative of an individual's ancestral origins (e.g. based on the birthplaces of recent ancestors, such as grandparents). We do not proscribe a general solution to this problem, and for the current defaults we use locations based on the approach taken in the source publications. A future feature will allow alternative location schemas to be used for the populations in a dataset.

We also envision a variety of future extensions to the GGV that would allow for further dissection of geographic structure in large-scale population genomic datasets. Providing an interactive means of browsing neighboring variant sites near a SNP of interest would offer a unique view into patterns of linkage disequilibrium around that focal SNP. This feature would be relevant to both medical geneticists conducting genome-wide association studies with interests in fine mapping as well as population geneticists interested in scanning the genome to detect signatures of positive selection. We imagine that incorporating a chromosomal browser such as jbrowse (SKINNER *et al.*, 2009) within the GGV would be greatly utilized by researchers and educators alike.

## Acknowledgements

Support for this work was provided by the National Institutes of Health via the Big Data to Knowledge initiative (1U01 CA198933-0) to JN and the National Institute of General Medicine under training grant award number T32GM007197 for JHM. The content is soley the responsiblity of the authors and does not necessarily reflect the official view of the National Institutes of Health. We acknowledge the Research Computer Center at the University of Chicago, especially H. Birali Runesha, Jeff Tharsen, Richard Williams, and Alex Mueller, for on-going support and extensions of the GGV browser. We also thank John Zekos for web server administration and support. The authors would also like to thank members of the Novembre Lab for supportive conversations.

## Appendix

### Example 1: Query by rsid

~~~
http://popgen.uchicago.edu/ggv_api/freq_table?data="1000genomes_phase3_table"&rsID=rs1834640
[
 {
    "alleles": ["A", "G"],
    "pos": ["-15.310139", "13.443182"],
    "pop": "GWD",
    "nobs": "226",
    "xobs": "17",
    "freqscale": 1,
    "freq": [0.0752212389381, 0.9247787610619],
    "chrom_pos": "15:48392165",
    "rawfreq": 0.0752212389381
  }, …
]
~~~

### Example 2: Query by chromosome position

~~~
http://popgen.uchicago.edu/ggv_api/freq_table?data="1000genomes_phase3_table"&chr=14&pos=37690093
[
  {
    "alleles": ["G", "A"],
    "pos": ["-15.310139", "13.443182"],
    "pop": "GWD",
    "nobs": "226",
    "xobs": "0",
    "freqscale": 0.01,
    "freq": [0.0, 1.0],
    "chrom_pos": "14:37690093",
    "rawfreq": 0.0
 }, …
]
~~~

### Example 3: Random query

~~~
http://popgen.uchicago.edu/ggv_api/freq_table?data="1000genomes_phase3_table"&random_snp=True
[
 {
    "alleles": ["T", "C"],
    "pos": ["-15.310139", "13.443182"],
    "pop": "GWD",
    "nobs": "226",
    "xobs": "0",
    "freqscale": 0.01,
    "freq": [0.0, 1.0],
    "chrom_pos": "5:42452893",
    "rawfreq": 0.0
 }, …
]
~~~

